# Induced Nanoscale Membrane Curvature Bypasses Clathrin’s Essential Endocytic Function

**DOI:** 10.1101/2021.07.15.452420

**Authors:** Robert C Cail, Cyna R Shirazinejad, David G Drubin

## Abstract

During membrane trafficking, flat membrane is rapidly remodeled to produce nanometer-scale vesicles. The mechanisms underlying this shape change are not completely understood, but coat proteins such as clathrin are implicated. Clathrin’s ability to bind to membranes of many different geometries casts uncertainty on its specific role in curvature generation and stabilization. Here, we used nanopatterning to produce substrates of ideal optical properties for live-cell imaging, with U-shaped features that bend a cell’s ventral plasma membrane into shapes characteristic of the energetically-unfavorable intermediate of clathrin-mediated endocytosis (CME). This induced plasma membrane curvature recruits the endocytic machinery, promoting productive endocytosis. Upon clathrin or AP2 disruption, CME sites on flat substrates are diminished. However, induced curvature rescues the localization, turnover and transferrin cargo uptake activities of CME sites after clathrin, but not AP2, disruption. These data establish that clathrin’s essential function during CME is to facilitate the evolution of membrane curvature rather than to scaffold CME protein recruitment.

**Summary:** Cail et al. demonstrate that induced nanoscale membrane curvature recruits endocytic sites and produces vesicles in cells lacking the coat protein clathrin, while the adaptor protein AP2 is still required, showing that clathrin’s essential function is to facilitate curvature development of the nascent vesicle.

## Introduction

Mammalian cell membrane shape is established and maintained in complex 3D environments, wherein nanoscale membrane bending is induced by both external environmental factors and intrinsic cellular processes (Chen et al., 1997). Features such as the extracellular matrix (ECM) and cell-cell junctions force the cell to adopt particular geometries, and intracellular processes like endocytic vesicle formation reshape the membrane from within (Suleiman et al., 2013; Liu et al., 2010; Kovtun et al., 2020). To generate vesicles, the cell must bend regions of its flat membrane into a spherical shape on the order of 100 nm across, an energetically costly process that requires passing through an unfavorable U-shaped transition state (Hassinger et al., 2017).

Clathrin-mediated endocytosis (CME), the best-studied form of extracellular cargo uptake, is a constitutive process that regulates a wide array of cellular functions including signaling, adhesion, and migration (Tan et al., 2018; Ezratty et al., 2009; Paul et al., 2015). To generate vesicles, adaptor proteins such as AP2 bind to cargoes, early curvature-generating proteins such as FCHo1/2, and the coat protein clathrin, marking a clathrin-coated pit (CCP) (Jackson et al., 2010; Henne et al., 2010). As the endocytic site matures, a feedback loop of curvature generation and stabilization occurs, with incorporation of more adaptors, cargoes, and curvature-generating proteins such as epsins, culminating in the production of a clathrin-coated vesicle and membrane scission catalyzed by the GTPase dynamin (Ehrlich et al., 2004; Ford et al., 2002; Grassart et al., 2014; Busch et al., 2015). CCPs localize preferentially to sites of nanoscale curvature in cultured cells, suggesting that induced plasma membrane geometry may play a physiological role in CCP nucleation and dynamics (Zhao et al., 2017).

Clathrin is essential for productive CME; upon clathrin disruption through siRNA or chemical inhibition, CCPs stall as protein aggregates with shallow curvature (Henrichsen et al., 2006; Von Kleist at al., 2011). However, clathrin can adopt distinct functional conformations including the spherical cage, flat plaque, and tubular lattice (Roth and Porter, 1964; Leyton-Puig et al., 2017; Elkhatib et al., 2017). While *in vitro* evidence has shown that clathrin preferentially binds to membranes of high curvature in a manner dependent on oligomeric assembly, *in situ* electron microscopy has emphasized the abundance of flat clathrin assemblies and provided evidence for a flat-to-curved transition of CCPs (Zeno et al., 2021; Sochacki et al., 2021; Avinoam et al., 2015). The observations that clathrin associates with membranes of different shapes and can assemble into different geometries have led to debate about clathrin’s activity during CME. What role does clathrin play in membrane budding, given its affinity for membranes of different curvatures? Answering this question promises to provide new insights into how productive CCPs are established and regulated in the cell, and into CME’s membrane curvature-generating mechanism. By creating nanofabricated ridges on a material with ideal properties for live-cell imaging that induce a range of nanoscale membrane curvatures, mimicking ECM fibers across the ventral cell membrane, we set out to answer these questions.

## Results

### Ormocomp nanoridges induce membrane curvature and recruit the endocytic machinery

To induce membrane curvature across nanometer-range size scales, we applied a low-cost, high-throughput nanofabrication method termed UV-Nanoimprint Lithography (UV-NIL) (Fig 1A, Fig S1A) (Bender et al., 2002). The principal advantage of UV-NIL is its ability to create sub-100-nm structures, below the limit for standard soft lithographic methods employed for cell biology studies, allowing the creation of structures on the scale of a macromolecular ensemble such as a CCP (Schmid and Michel, 2000). We selected the organic/inorganic hybrid material Ormocomp as our substrate because of the near-exact match of its refractive index to that of borosilicate glass, allowing us to conduct Total Internal Reflection Fluorescence (TIRF) microscopy, a standard imaging method in the CME field (Fig S1B) (Gissibl et al., 2017; Merrifield et al., 2002).

**Fig 1.**
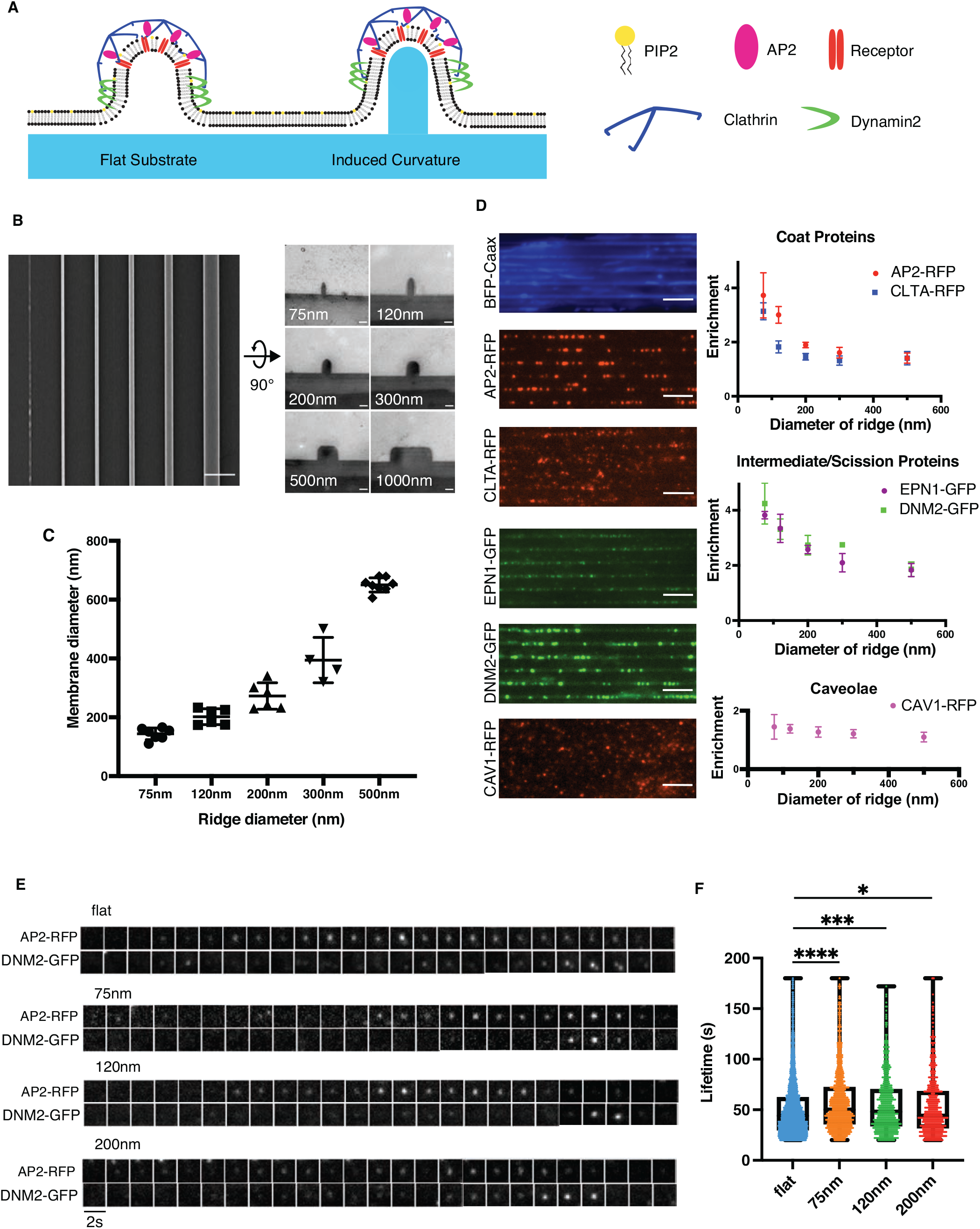
**(A)** Model of Ormocomp-induced stabilization of endocytic sites, mimicking the U- shaped intermediate stage of CME. **(B)** Right: En-face SEM image of nanoridge structures. Scale bar: 2 µm. Left: thin-section TEM demonstrating the shape and scale of protrusions on Ormocomp ridges. Scale bar: 100nm. **(C)** The diameter of membrane induced by ridges of various diameter. N ≥ 3 ridges from 3 cells. Mean +/- SD. **(D)** Left: Survey of endocytic proteins demonstrating enrichment of endocytic sites on sites on 75nm nanoridges (scale bar 4 µm). Quantification of enrichment score demonstrates the increase in endocytic localization as a function of curvature for early, intermediate, and late proteins, as well as a marker of caveolae. Mean +/- SD. N ≥ 4 cells per fluorescent protein. **(E)** Representative montages of AP2RFP/ DNM2GFP endocytic sites from flat, 75nm, 120nm, and 200nm substrates, showing canonical profile of AP2RFP and DNM2GFP marking productive endocytosis. **(F)** Lifetimes of endocytic sites as a function of curvature, showing increased lifetime for productive CME at smallest pillars with decrease as pillar diameter increases. P values: **** > 0.0001, *** > 0.001, * > 0.05, Student’s T test. Mean with interquartile range. N ≥ 1,000 tracks per ridge size.

To produce substrates through UV-NIL, we first created a mold through electron-beam lithography and reactive-ion etching on a silicon wafer (Fig S1A) (Fischer and Chou, 1992). This mold features mm-long, cylindrical invaginations spaced at a pitch of 2 µm, with widths of 75, 120, 200, 300, 500, and 1000 nm (Fig 1B). The depth of an invagination is 200 nm for 75 nm ridges, 250 nm for 120 nm ridges, and 300 nm for larger sizes. This mold served as the basis for Ormocomp substrate production. The resulting Ormocomp substrates bear the inverse geometry of the mold and therefore have cylindrical protrusions, which we term nanoridges.

Upon exposure to intense UV irradiation, Ormocomp polymerizes through polyacrylate chemistry. Ormocomp substrate manufacturing requires only ∼15 minutes per substrate, making replicated experiments and wider screening of conditions more practical than with more standard, slower nanofabrication processes.

To determine how the nanoridges affect plasma membrane shape, we seeded MDA-MB-231 cells on nanoridge substrates and performed thin-section TEM of substrate-grown cells to measure the diameter of the membrane invaginations induced by each nanoridge (Fig S1C). By measuring the diameter of multiple ridge-membrane interfaces across >3 cells per ridge size, we found that the nanoridges induce consistent membrane diameter as a function of ridge size (Fig 1C). The smallest nanoridge, 75 nm, induced a membrane diameter of ∼142 nm, approximately 50% larger than the diameter of a fully-formed vesicle, with a linear increase up to the 500 nm ridge, which induced ∼650 nm diameter curvature. The shape of the membrane bent around nanoridges was similar to the shapes seen in MDA-MB-231 cell membranes wrapped around collagen fibers, which were shown to induce clathrin structures known as Tubular Clathrin/AP2 Lattices (TCALs) (Elkhatib et al., 2017). Thus, Ormocomp nanoridges provide a robust method to create regular, reproducible membrane curvature and to de-couple curvature generation from cell-intrinsic processes such as vesicle formation.

To determine the effect of the diameter of induced membrane curvature on the endocytic machinery, we seeded MDA-MB-231 cells, genome-edited to express fluorescent fusion proteins under endogenous regulatory control, on nanoridge substrates and imaged them on a TIRF microscope (Fig 1D, Fig S1D) (Doyon et al., 2011). Imaging of plasma membrane markers showed that the entirety of the ventral plasma membrane is illuminated by the TIRF evanescent field (Fig 1D, Fig S1E). The endocytic proteins demonstrated a striking preference for the higher-curvature ridges (e.g., 75, 120, and 200 nm) (Fig 1D), as reported previously for cells grown on quartz substrates (Zhao et al., 2017). This observation was true of early-arriving endocytic proteins AP2 and clathrin light chain A, as well as the intermediate curvature-generating protein epsin1 and late-peaking scission GTPase dynamin2. The protein caveolin1, which is a marker of caveolae and not a component of CME sites, demonstrated no preferred binding to membranes of high curvature (Rothberg et al., 1992).

Because Ormocomp nano ridges promised better optical properties than the previously used quartz substrates and they could be produced with higher throughput, enabling more conditions to be analyzed, and because they had not previously been used for studies of CME, we next performed an image-based analysis of CME proteins associated with these structures. To quantify the enrichment of endocytic proteins on nanoridges with high curvature, we developed automated image processing tools. Using the bright field image of a cell on a substrate, we employed a line detection algorithm to create a mask that discriminates between on-and-off ridge sections of the cell. We next employed a Differences-of-Gaussians (DoG) algorithm to detect and localize fluorescent puncta (Tinevez et al., 2017). We corrected for increased cell membrane presence on nano-ridges by measuring the on-ridge increase in fluorescence of cell membrane markers BFP-Caax and CellMask Orange, which were in good agreement (Fig S1E). By measuring the relative abundance of endocytic proteins on the ridge vs flat membrane and dividing by the corrected area of the mask, we determined an enrichment score, which is a relative increase in the likelihood of endocytic protein localization as a function of curvature. An enrichment score of 1 corresponds to no increase in localization, while scores greater than 1 indicate an increase in the number of fluorescent puncta compared with flat membrane.

The enrichment score demonstrates the strong preference of the endocytic machinery for regions of high plasma membrane curvature (Fig 1D). At the 75-nanometer ridge, AP2-RFP shows a nearly 4-fold increase in puncta localization relative to flat membranes. Epsin1-GFP and dynamin2-GFP both show greater than a 4-fold enrichment. Swapping the fluorophores on AP2 and dynamin2 to GFP and RFP, respectively, resulted in little difference in enrichment (Fig S1F). These enrichments decrease with increasing diameter, until at the 500-nm ridge there is essentially no detectable enrichment. Clathrin light chain A, in contrast with other endocytic proteins, demonstrates a >3-fold enrichment at the smallest substrate size but precipitously decreases its enrichment as ridge size increases. This reduced enrichment may be because of the detection of clathrin at other trafficking events within the cell body such as endosomes and the *trans*-Golgi network, above the ventral plasma membrane but present in the TIRF field (Stoorvogel at al., 1996; Daboussi et al., 2012). Caveolin1-RFP showed no change in enrichment score as a function of curvature, despite caveolar endocytosis being another form of endocytic uptake. These data demonstrate that CME proteins—but not necessarily other endocytic processes—robustly reorganize across the cell in response to induced membrane curvature, with a strong preferential localization at highest induced positive curvature.

We next wondered whether the puncta of endocytic proteins present on nanoridge- induced membrane curvature represent productive CCPs, TCALs, or ectopic protein localization. We imaged double-genome-edited MDA-MB-231 cells expressing AP2-RFP and DNM2-GFP at 0.5 Hertz on a range of ridge sizes. Montages and kymographs from two-color movies on flat and curved Ormocomp substrates demonstrated the characteristic track of gradual AP2-RFP fluorescence increase accompanied by a burst of DNM2-GFP at the end of each track, a hallmark of productive endocytosis (Fig 1E, Fig S2A) (Doyon et al., 2011). Through TEM imaging, we found the canonical spiked coat of clathrin on nanoridges and CCPs budding from them (Fig S2F). Using automated site detection and tracking and binning based on nanoridge size, we measured the lifetime of valid endocytic sites and found that the AP2 lifetime on flat substrates was 50.9+/-28s, very similar to previously reported values for CME sites on glass (Fig 1F) (Aguet et al., 2013; Hong et al., 2015). Endocytic sites on induced curvature had slightly lengthened lifetimes, from 58.4+/-30s for 75 nm ridges to 54.8+/-28s for 120 nm ridges and 53.5+/-29s for 200nm ridges. At diameters > 200 nm, endocytic dynamics were unchanged by nanoridges (Fig S2D). The nanoridges did not change the percentages of transient, broken, incomplete, or persistent CCPs, and did not change the bulk rate of endocytosis as measured by fluorescent transferrin uptake assay (Fig S2E, Fig S3A-B). These data indicate that localized curvature preferentially induces endocytic sites without altering overall endocytic profile.

TEM imaging of the 1000-nanometer substrate revealed a flat top with curved sides (Fig 1B); cells grown on this size of ridge bear membrane curvature along the side of the ridge and flat membrane along the top of the ridge (Fig S3C). The diameter of the approximate semi-circle on this curved edge of the ridge is ∼210 nanometers, similar to that of the 120-nm nanoridge (Fig S3D). Consistent with the conclusion that substrate-induced membrane curvature recruits the endocytic machinery, AP2-RFP preferentially appears at the edges of the 1000-nm ridges (Fig S3E). Indeed, a CCP can be seen by TEM protruding from the high-curvature edge (Fig S3C).

### Induced curvature rescues endocytic site formation upon clathrin knockdown

Because membrane invaginations formed by U-shaped nanoridges are decoupled from natural endocytic site formation, we wondered whether induced curvature might bypass functions of endocytic proteins such as clathrin. Upon siRNA treatment, clathrin heavy chain (*CLTC*) expression was reduced by approximately 95%, as measured by western blot (Fig 2A). Previously, it was established that CME in mammalian cells is inhibited when clathrin is knocked down, leading to unstable nascent endocytic sites that stall as flat patches (Henrichsen et al., 2003; Henrichsen at al., 2006). Consistently, compared to sites seen in cells treated with control siRNA, clathrin knockdown on flat substrates leads to dimmer AP2-RFP sites with markedly reduced DNM2-GFP puncta intensity and less overlap with AP2-RFP, indicating that true endocytic sites are not formed (Fig 2Bi).

**Fig 2.**
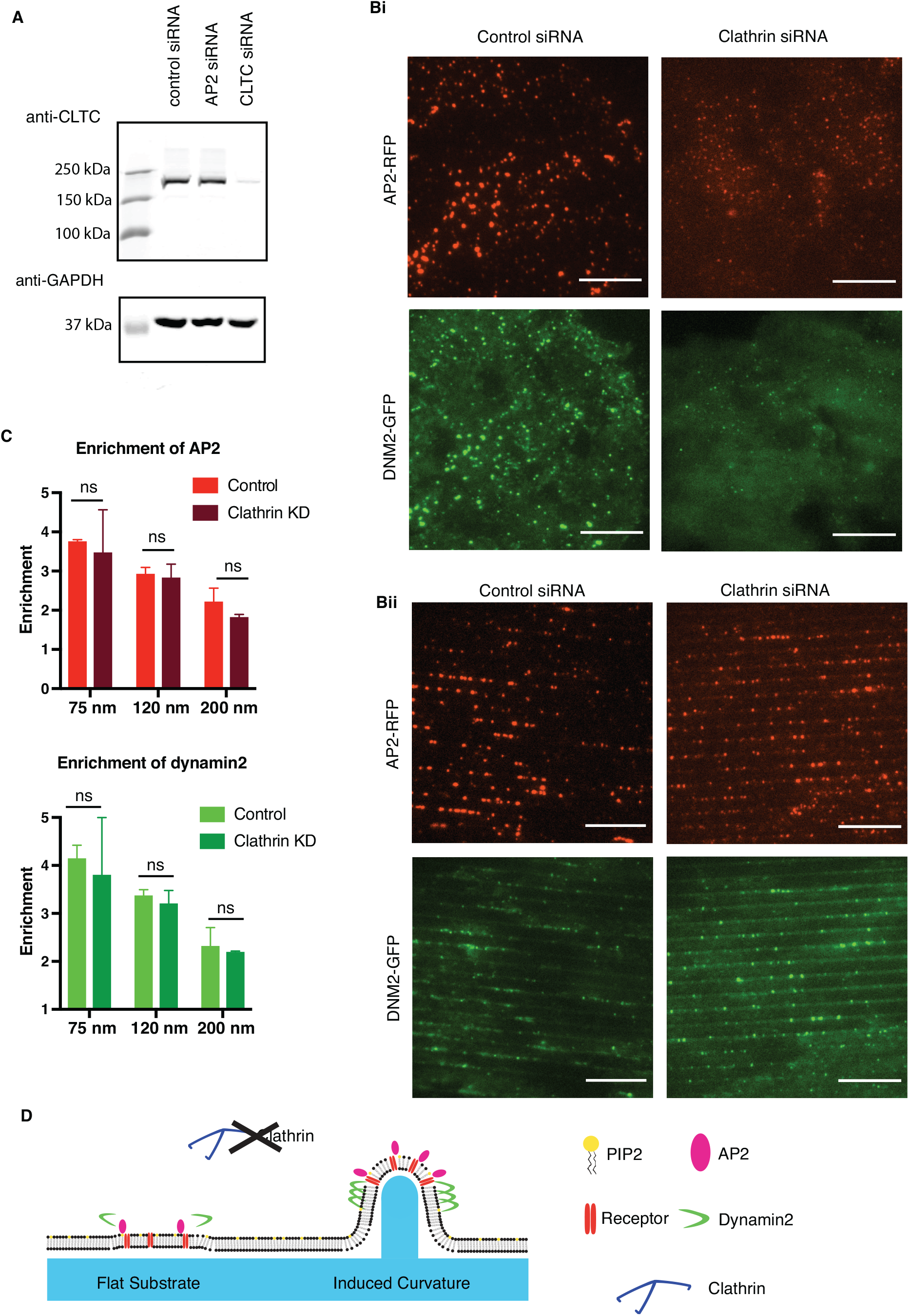
**(A)** Western blot against clathrin heavy chain from control siRNA, *AP2µ* siRNA, and *CLTC* (clathrin) siRNA cells demonstrating the knockdown efficiency of clathrin knockdown. Western blot against GAPDH is the loading control. **(Bi)** Representative images of AP2RFP and DNM2GFP from control and clathrin siRNA cells on flat Ormocomp. **(Bii)** Representative images of AP2RFP and DNM2GFP from control and clathrin siRNA cells on 75nm substrate, showing a rescue of number and intensity of AP2RFP and DNM2GFP puncta. Scale bars: 10 µm. **(C)** Enrichment score of AP2RFP and DNM2GFP across substrate size. P-values > 0.05, Multiple T tests. Mean +/- SD. N = 3 cells per condition. **(D)** Model of endocytic site stabilization lacking clathrin, but with or without induced curvature to stabilize the endocytic site.

However, when clathrin knockdown cells were grown on 75-200 nm nanoridges, there was at the sites of highest curvature a striking enrichment in AP2-RFP puncta and an increase in puncta intensity and apparent site size, as well as robust enrichment of DNM-2GFP puncta (Fig 2Bii). Thin-section TEM imaging revealed that the diameter of the membrane on nanoridges was unaffected by clathrin knockdown (Fig S4A-B). The enrichment score for AP2-RFP and DNM2- GFP was unchanged by clathrin siRNA, indicating that these early- and late-arriving CME proteins localize to sites of induced membrane curvature independent of clathrin (Fig 2C). These data support a model of endocytic site formation in which, in the absence of clathrin, induced curvature is sufficient to stabilize the endocytic site and localize both early and late endocytic machinery (Fig 2D).

### Induced curvature does not rescue endocytic site formation upon AP2 knockdown

To test whether formation of curvature-dependent CME sites requires the adaptor protein AP2, or whether the CME sites can form with curvature but without receptor clustering, we knocked down AP2 expression by siRNA and imaged cells on flat and curved substrates. Upon siRNA treatment, AP2 expression was depleted by >90%, as demonstrated by western blot and by a decrease in AP2 fluorescence in AP2RFP-tagged cells (Fig 3A, Fig S4C). AP2 disruption leads to a marked decrease in CLTA-RFP and DNM2-GFP puncta on flat substrates. This decrease in endocytic proteins is not rescued by induced curvature (Fig 3B). AP2-depleted cells on a flat substrate showed 3-fold decreased density of DNM2-GFP puncta, and 2-fold decreased density of CLTA-RFP puncta (Fig 3C). Despite the diameter of induced membrane curvature being unchanged by AP2 siRNA, the enrichment score for CLTA-RFP decreased significantly upon AP2 knockdown, from approximately 2.2 to 1.1 on 75nm substrates; this decrease in enrichment score is present across substrate sizes, indicating that clathrin is curvature-insensitive after AP2 disruption (Fig 3D). The remaining DNM2-GFP puncta, however, were still 2-fold enriched on sites of high curvature, which is a more modest enrichment than in control cells (Fig 3D). These data indicate that AP2 is an essential component of the endocytic machinery irrespective of induced curvature, and that curvature stabilization is downstream of receptor- adaptor interactions.

**Fig 3.**
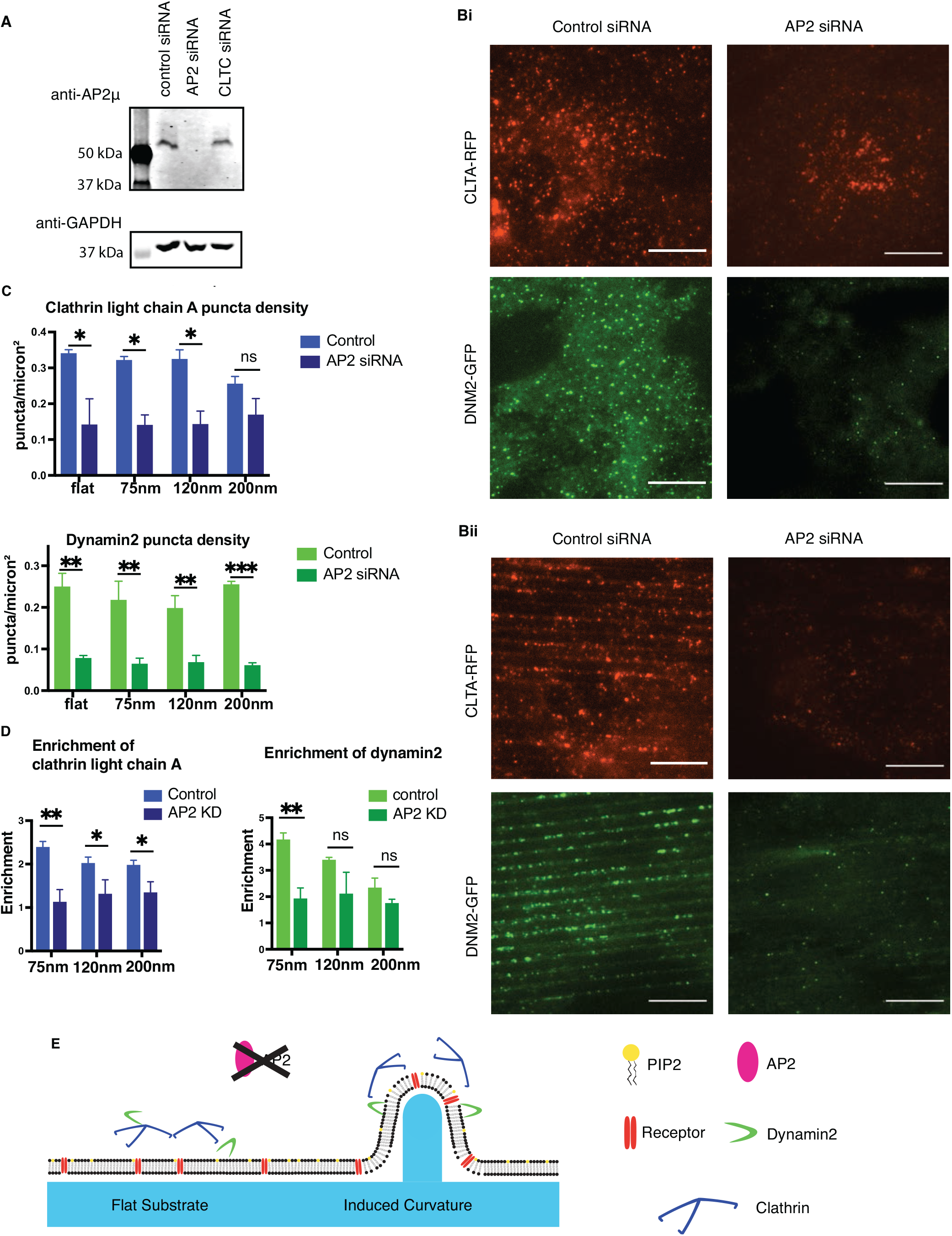
**(A)** Western blot against AP2 µ-subunit from siRNA, *AP2µ* siRNA, and *CLTC* siRNA cells demonstrating the knockdown efficiency of AP2 knockdown. Western blot against GAPDH is the loading control. **(Bi)** Representative images of CLTARFP and DNM2GFP from control and AP2 siRNA cells on flat substrate showing reduction in endocytic site formation in response to AP2 disruption. **(Bii)** Representative images of CLTARFP and DNM2GFP from control and AP2 siRNA cells on 75nm nanoridges. Scale bars: 10 µm. **(C)** Quantification of number of CLTARFP and DNM2GFP puncta across substrate size with control or AP2 siRNA. P values: ** > 0.01, * > 0.05, ns > 0.05 from Multiple T test. Mean +/- SD. N = 3 cells per condition. **(D)** Enrichment score of CLTARFP and DNM2GFP on substrates with control or AP2 siRNA. P values: ** > 0.01, * > 0.05, ns > 0.05 from Multiple T test. Mean +/- SD. N = 3 cells per condition. **(E)** Model of endocytic sites with or without induced curvature in response to AP2 knockdown, showing lack of stabilization of endocytic proteins without central AP2 hub.

### Endocytic sites on curved substrates show characteristic fluorescence dynamics profiles even without clathrin

Once we established that AP2 and dynamin2 localization in clathrin knockdown cells could be rescued by induced membrane curvature, we wondered whether these rescued sites turn over with characteristic dynamics even without the clathrin coat, and what their fluorescence profile over time would be. To determine the kinetic profile for these sites over time, we recorded time-lapse TIRF microscopy of AP2/dynamin2 double-labeled cells after clathrin knockdown and tracked the endocytic sites over time. We found that for cells treated with control siRNA, the sites of AP2/dynamin2 colocalization appeared as canonical endocytic sites, with a gradual increase in AP2 accompanied by a burst of dynamin2 to indicate scission (Fig 4A). Upon clathrin knockdown on flat substrates, the AP2 puncta were dimmer, small and unstable, with reduced dynamin2 fluorescence. However, with substrate-stabilized 75 nm membrane curvature, the AP2 puncta became much larger and more stable. Additionally, there was a burst of dynamin2 at a late stage of AP2 lifetime, followed by a decrease in AP2 fluorescence signal, hallmarks of CME vesicle scission. We found that the average lifetime of these tracks in clathrin knockdown cells (68+/-33s) was significantly increased over what was observed in control siRNA cells (50+/-26s), indicating that despite the normal appearance of the sites, their turnover is affected by clathrin knockdown (Fig 4C).

**Fig 4.**
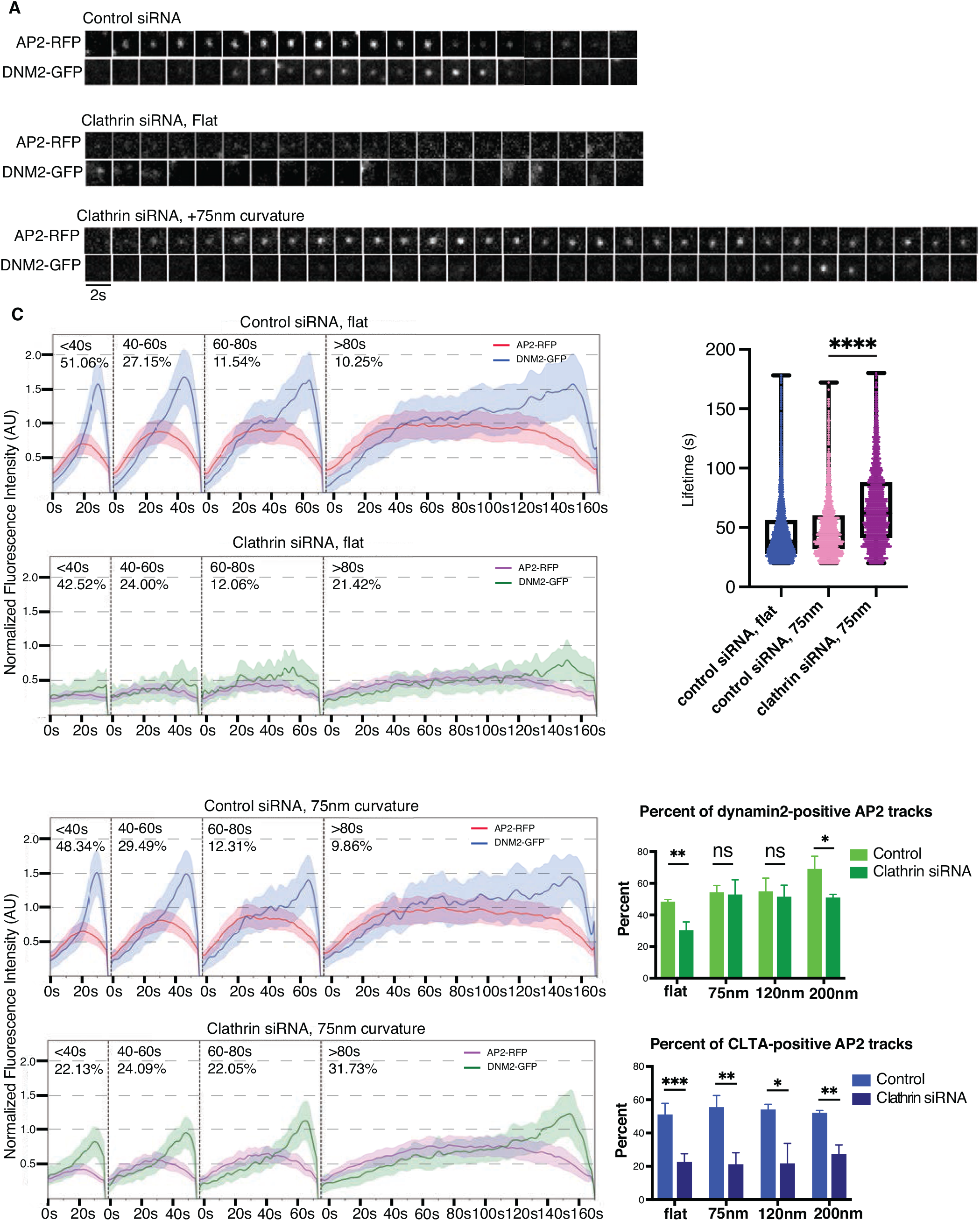
**(A)** Representative montages of AP2RFP/DNM2GFP endocytic sites from control siRNA and clathrin siRNA with or without 75nm of induced curvature. **(Bi)** Normalized fluorescence intensity profiles for endocytic sites binned by lifetime from flat substrates, with or without clathrin knockdown. **(Bii)** Normalized fluorescence intensity profiles for endocytic sites binned by lifetime from substrates with 75nm nanoridges, with or without clathrin knockdown, demonstrating increase in AP2/DNM2 fluorescence intensity and late peak of DNM2. N ≥ 800 tracks per condition. **(C)** Lifetimes of endocytic sites from flat and 75nm control siRNA endocytic sites and 75nm clathrin siRNA sites, showing an increase in lifetime with clathrin disruption. P-values: **** > 0.0001, Student’s T test. Mean with interquartile range. N ≥ 800 tracks per condition. **(D)** Overlap of AP2/DNM2 or AP2/CLTA with control or clathrin siRNA as a function of substrate size. P-values: P values: *** > 0.001, ** > 0.01, * > 0.05, ns > 0.05 from Multiple T test. Mean +/- SD. N = 3 cells per condition.

After binning the AP2/dynamin2 positive tracks according to lifetime, we found that, for cells grown on flat substrates, clathrin knockdown results in a fluorescence signal decrease to only ∼35% for AP2/dynamin2 tracks relative to control siRNA-treated cells (Fig 4Bi). The remaining dynamin2 signal showed a nearly symmetrical, noisy distribution over time, rather than a late-peaking recruitment characteristic of cells treated with control siRNA. However, tracks for cells grown on 75nm substrates demonstrated a rescue of fluorescence intensity to ∼70% of control, and the dynamin2 signal demonstrated late peaks characteristic of productive vesicle formation (Fig 4Bii).

In addition to the large decrease in intensity of AP2-RFP puncta after clathrin heavy chain knockdown, there was a significant decrease in detected overlap between AP2-RFP and DNM2- GFP with automated track detection (Aguet et al., 2013; Burckhardt et al., 2021). The percentage of AP2-RFP detections that overlap with DNM2-GFP decreased from 49% to 29% (Fig 4D).

Quantification of the overlap between AP2-RFP detections with DNM2-GFP on 75nm ridges after clathrin knockdown demonstrates a dramatic rescue in overlap between the two fluorophores, to 52%, matching the overlap with control siRNA. This increase in overlap persists across nanoridge size up to 200nm. In contrast, in cells knocked down for clathrin heavy chain, overlap between AP2-GFP detections and clathrin light chain A (CLTA-RFP) was markedly decreased from 51% to 22%, and this decrease in fluorescence overlap is not changed by induced curvature, evidence that the rescue of the endocytic machinery is not provided by residual clathrin heavy chain.

### Curvature-rescued CME sites take up endocytic cargoes

Once we saw that curvature-rescued CME sites turned over with similar fluorescence profiles to control CME sites, we next sought to determine whether these sites could also take up cargo in the absence of clathrin. To test this, we performed transferrin uptake assays with Alexa- Fluor 647-labeled transferrin (tfn-AF647). After serum starvation, upon exposure to media containing tfn-AF647, control siRNA-treated cells demonstrated hundreds of trafficked puncta of tfn-AF647 in the cell body (Fig 5A). Clathrin siRNA cells on flat substrate demonstrated a largely reduced number of weakly-fluorescent puncta apparently associated with the cell surface (see below) from transferrin molecules binding to their receptor and clustering but failing to achieve internalization. With 75nm-curvature, however, clathrin heavy chain siRNA cells demonstrated a marked increase in number and intensity of tfn-AF657 puncta. To quantify the uptake of fluorescent cargo, we drew regions-of-interest around individual cells and measured the mean pixel intensity across 10 cells per condition. We found the mean pixel intensity of control cells to be ∼1200+/-500 AU (Fig 5B). Clathrin knockdown cells on flat substrate had a mean pixel intensity of 69+/-22AU, while clathrin knockdown cells with 75nm of induced membrane curvature demonstrated a >3-fold increase in mean pixel intensity to 223+/-73 AU. The intensity of curvature-rescued cells was >4-fold less than for control cells, which is consistent with 1) exogenous curvature only partially rescuing endocytosis and 2) only ∼⅓ of the cell, the ventral membrane contacting nanoridges, having induced membrane curvature.

**Fig 5.**
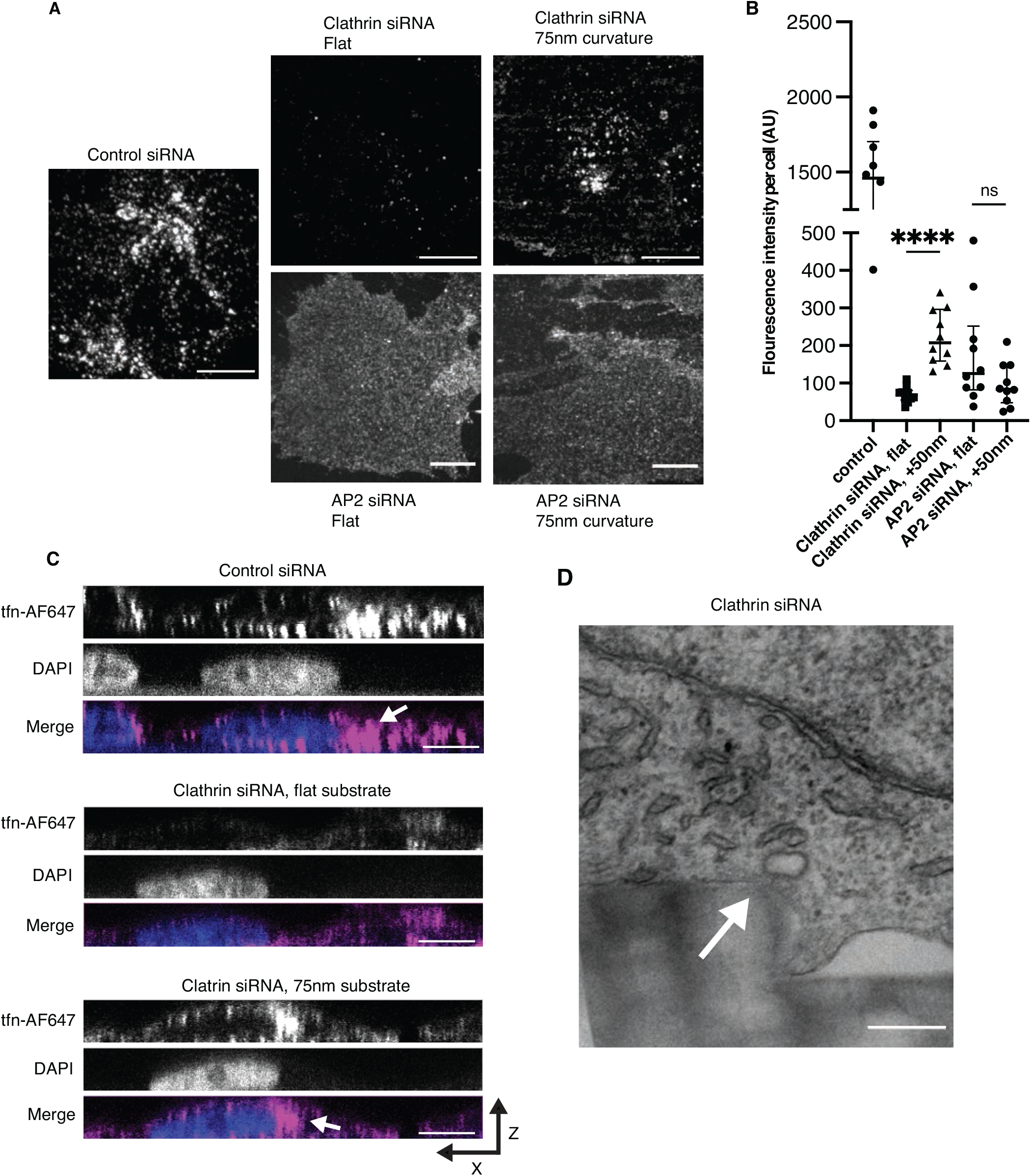
**(A)** Representative images of AlexaFluor-647 tagged transferrin (tfn-AF647) in control siRNA, clathrin siRNA, and *AP2µ* siRNA cells with or without induced curvature. **(B)** Quantification of mean fluorescence intensity of cells from **(A)**. P-values: P values: **** > 0.0001, ns > 0.05 from Student’s T test. Mean +/- SD. N = 10 cells per condition. **(C)** X-Z projections of cells with control or clathrin siRNA cells with or without induced curvature, with white arrow marking tfn-AF647 next to DAPI stain showing the nucleus. **(D)** TEM image from clathrin siRNA cell with induced curvature, with white arrow marking a putative vesicle that lacks a clear clathrin coat but forms on the edge of a site of high induced curvature.

To determine where the puncta of tfn-AF647 are located within the cell, we created X-Z projections of confocal microscopy images (Fig 5C). Control cells demonstrated perinuclear localization of tfn-AF647, indicating trafficking to an endosomal compartment after uptake. In contrast, clathrin siRNA cells on flat substrate showed only membrane-associated tfn-AF647 puncta. However, with induced curvature, we observed perinuclear tfn-AF647 puncta in clathrin knockdown cells, demonstrating that not only is there an increase in clustering of cargo with curvature but that these clustered sites are potentiated for uptake even without the requisite coat. Through TEM imaging of clathrin knockdown cells, we found associated with the plasma membrane on regions of induced membrane curvature, membrane-enclosed structures that lack a clear clathrin coat and have an elliptical shape; these might represent vesicles produced through curvature-induced rescue of clathrin disruption near the time of scission (Fig 5D).

To test whether curvature can rescue cargo uptake in the absence of AP2, we repeated the transferrin uptake assay in the context of AP2 knockdown (Figure 5A). AP2 knockdown leads to membrane-associated tfn-AF647 signal, as the transferrin binds to its receptor on the plasma membrane but cannot cluster or be taken into the cell. AP2-knockdown cells grown on 75- nanometer ridges demonstrated tfn-AF647 evenly on the plasma membrane, and no puncta were visible over the diffuse membrane signal. No statistically significant change in average pixel intensity in AP2-knockdown cells was observed in response to induced membrane curvature (Fig 5B). This result indicates that transferrin and/or its receptor itself are not detectably curvature sensitive, and that CME protein preference for high membrane curvature is downstream of AP2 function.

## Discussion

In this study, to gain insights into the feedback between membrane geometry and biochemical activity, we created induced nanoscale curvature in live cells using a novel substrate system with optimal optical properties for live-cell imaging. An attractive hypothesis for how membrane curvature development is controlled during vesicle formation is that the membrane geometry itself acts as a signal to control biochemical reaction rates such as Bin-Amphiphysin-RVS domain protein recruitment, receptor clustering or lipid phosphatase activity (Liu et al., 2009). Indeed, it has been shown *in vitro* that high curvature and specific lipid characteristics coordinate to induce actin polymerization, as would occur at a late stage of endocytosis (Daste et al., 2017). Membrane geometry and biochemical reaction rates might feed back on each other across stages of CME as a flat membrane evolves into a fully mature, spherical vesicle.

The enrichment score used in this paper gives useful mechanistic insight into the stepwise progression of vesicle formation and the characteristic affinities of proteins for curvature; for instance, we hypothesize that caveolin1, with its low enrichment on curved membrane, lacks affinity for high curvature because of its localization on stable membrane tubules, rather than requiring a flat-to-curved transition for vesiculation. The decrease in clathrin’s enrichment after AP2 knockdown demonstrates that clathrin relies on AP2 and other proteins to stabilize an endocytic site. Interestingly, in MDA-MB-231 cells used in this study, dynamin2 still bore some curvature enrichment, although significantly lower than in control cells, even after AP2 knockdown eliminated essentially all productive CME sites. This enrichment may be due to some innate curvature preference for dynamin2, or it may be due to dynamin2’s roles in other curvature-sensitive endocytic pathways (Schafer et al., 2002; Henley et al., 1998).

Clathrin’s role during CME has been a topic of debate since the discovery through electron microscopy some 40 years ago of its heterogeneous conformations on the plasma membrane (Heuser and Evans, 1980). While many CME proteins have been shown to sense curvature on their own in purified systems, clathrin’s ability to respond to curvature *in vitro* depends on its oligomeric assembly into a macromolecular ensemble and is greatly enhanced by other proteins present during CME (Busch et al., 2015; Zeno et al., 2018; Zeno et al., 2021). One proposed model for endocytic curvature formation is that clathrin does not itself generate curvature, but acts as a Brownian ratchet, leveraging subunit exchange as the pit grows to irreversibly stabilize the membrane curvature generated by thermal fluctuations and membrane-bending proteins (Sochacki and Taraska, 2018; Avinoam et al., 2015). Consistent with this model, we find that our artificially induced nanoscale curvature itself is sufficient to produce vesicular uptake in the absence of a clathrin coat. This curvature-induced endocytosis may be relevant to other trafficking events such as compensatory endocytosis, where coat proteins might be absent, or in divergent endocytosis such as viral uptake, as viral particles are orders of magnitude stiffer than the human plasma membrane and are often larger than canonical CCP cargoes (Sokac at al., 2003; Mateu, 2012; Ehrlich et al, 2004).

The high-throughput UV-NIL approach described here to nanofabricate substrates from Ormocomp, uniquely enables both precise control of membrane curvature and state-of-the-art imaging. When coupled with quantitative cellular assays, this approach will allow in vivo studies of many other curvature-sensitive proteins so how 3D nano-geometry affects biochemical reaction rates underlying diverse cellular processes can be determined.

## Materials and Methods

### Substrate Nanofabrication

Nanoridge substrates were made from Ormocomp (Microresist Technologies) adhered to Fisher #1 25mm coverslips (12-545-86). Coverslips were prepared by cleaning with isopropyl alcohol followed by spin-coating at 2000 RPM for 45 seconds covered with approximately 50 µL of hexamethyldisilazane (Sigma-Aldrich, 440191) and 5-minute curing at 180 degrees.

Ormocomp was shaped using a silicon wafer mold, and polymerized by exposure to 110 seconds of UV at 14 mW on OAI Mask Aligner model 204. Silicon molds were made using EBL (Vistec VB300) on ZEP520A resist followed by Inductively-Coupled Plasma Reactive Ion Etching using Sulfur Hexafluoride (2%) and molecular Oxygen (98%) at -150 deg in liquid nitrogen-cooled chambers (Oxford Plasmalab RIE). ZEP520A was removed from silicon using acetone/IPA washes. Hydrophobicity of silicon mold was maintained by gas-phase trichloro- perfluoroctylsilane deposition (Sigma-Aldritch 448931-10G) for 30 minutes in vacuum chamber followed by curing at 150 deg for 10 minutes. Polymerized Ormocomp substrates were stored at room temp for up to two weeks prior to use, and irradiated with UV light for 12 minutes to sterilize prior to use in cell culture. Ormocomp substrate shape was determined by SEM imaging: Ormocomp was sputter-coated in titanium to a thickness of 1-2 nanometers and imaged on Hitachi S-5000 SEM.

### Cell Culture and transfection

MDA-MB-231 cells were genome-edited^19^ to express fluorescent fusion proteins. Cells were grown in DMEM/f12 (Life Technologies 10565-018) with 10% (v/v) FBS (Seradigm 89510-186) and Penicillin/Streptomycin (Life Technologies 15140-122) to no more than 20 passages. Cells were plated directly onto sterilized Ormocomp substrates at 70% confluence (approximately 8 x 10^5 cells per well) 18-24 hours prior to experiments or imaging.

Cells were prepared for TEM by fixation in 2% gluteraldehyde in cytoskeleton stabilization buffer (100 mM methyl ester sulfonate, 150 mM NaCl, 5 mM EGTA, 5 mM MgCl2, and 5 mM glucose in ddH2O, pH 6.8), rinsed 3x in PBS pH 7.4, stained in 1% Osmium tetroxide and 1.6% Potassium ferricyanide in PBS pH 7.4, and rinsed 3x in PBS pH 7.4. Cells were dehydrated in 7- minute washes in 30%, 50%, 70%, 95%, 100%, and 100% ice-cold ethanol and infiltrated in epon-araldite resin. Resin was polymerized in a 60-degree oven for 24 hours, then 70 nm sections were cut with an Ultracut E (Leica) and collected onto formvar-coated 50 mesh copper grids. Grids were post-stained with 2% uranyl acetate followed by Reynold’s lead citrate for 5 minutes each. Sections were imaged using a Tecnai 12 120kV TEM (FEI) and data recorded using an UltraScan 1000 with Digital Micrograph 3 software (Gatan Inc.). Membrane diameter measurements were made using ImageJ.

BFP-Caax in mammalian expression vector (pTagBFP2-C1), a generous gift from the Michael Rape lab (UC Berkeley), was transfected into cells on Ormocomp substrates at 70% confluence using lipofectamine 3000 (Invitrogen L3000008) according to manufacturer protocols ^(^https://tools.thermofisher.com/content/sfs/manuals/lipofectamine3000_protocol.pdf). Cells were imaged 18-24 hours after transfection. CellMask Orange (Invitrogen C10045) was diluted 1000x in media prior to staining; staining was performed for 10 minutes at 37 degrees, followed by two washes in fresh 37-degree media prior to imaging.

Dharmacon smartpool siRNAs (L-004001-01-0005, L-008170-00-0005, and D-001810-10-05) were used to knockdown Clathrin Heavy Chain or AP2Mu1, or a non-targeting control was used, respectively. Cells were transfected using lipofectamine 3000 according to manufacturer protocols (https://tools.thermofisher.com/content/sfs/manuals/lipofectamine3000_protocol.pdf), using an siRNA final concentration of 20 nM. Cells were incubated for 72 hours post-transfection prior to imaging. 18-24 hours prior to imaging or experiment, cells were split. 50% of cells were seeded on Ormocomp substrates. The remaining cells were re-plated, grown until imaging of Ormocomp substrates, then harvested for western blot analysis to determine knockdown efficiency. For western blots, cells were washed on ice in cold DPBS (Gibco 14190-144), lysed on ice using pre-chilled cell scrapers (Corning 3008) in lysis buffer (150 mM NaCl, 1% NP-40, 50mM Tris pH 8.0, 1x Roche protease inhibitor tablet (11697498001)), with immediate addition of 4x Laemmli buffer and 0.5% (v/v) beta-mercaptoethanol. Samples were thoroughly mixed by scraping tubes along a tube rack, boiled for 5 minutes, then centrifuged at 17,900 x g for 5 min before running at 120V on 10% SDS-polyacrylamide gel. Gels were transferred to nitrocellulose membrane (GE Healthcare 10600006) for 90 minutes at 40V in 4-degree transfer buffer (25 mM Tris base, 192 mM glycine, 20% (v/v) methanol), then blocked for 1 hour at room temperature in 5% milk in TBSt (10 mM Tris, 0.09% (w/v) NaCl, 0.05% (v/v) tween-20, pH 7.5). Membranes were probed with clathrin heavy chain antibody (Abcam ab1679) at 1:500 dilution in TBSt with 5% milk, AP2M1 antibody (Abcam ab75995) at 1:1000 dilution in TBSt with 5% milk, or GAPDH antibody (Abcam ab9485) at 1:2000 dilution in TBSt, all for 1 hour at room temperature, followed by 4 x 10- minute wash in TBSt. Secondary antibodies (LICOR, 926-32213) were blotted at 1:2500 dilution in TBSt for 1 hour at room temperature, followed by 4 x 10-minute wash in TBSt and immediate imaging on LICOR Odyssey CLx.

Transferrin uptake assays were performed by serum-starving cells in DMEM/f12 with 0.5% (w/v) BSA (Sigma Life Sciences A9647) for 30 minutes followed by adding fresh DMEM/f12/ BSA with alexa-fluor 647-labeled transferrin at a concentration of 10 micrograms/milliliter (Jackson Immunoresesarch 009-600-050). Cells were exposed to transferrin for 2 minutes followed by 2 minute chase in fresh DMEM/f12/BSA. Cells were then washed 1x with ice-cold DPBS. Cells were fixed in 4% paraformaldehyde (Electron Microscopy Solutions) in Tris/ potassium chloride cytoskeleton buffer (10mM MES, 150mM NaCl, 5mM EGTA, 5mM glucose, 5mM MgCl2, 0.005% Sodium Azide, pH 6.1) for 20 minutes at room temperature, followed by 3 x 5-minute washes in 50mM ammonium chloride in cytoskeleton buffer, and mounted on microscope slide in fluorescence mounting medium (Vectashield H-2000) and either imaged immediately on a confocal microscope or stored at 4 degrees overnight until confocal imaging.

### Microscopy

TIRF microscopy was carried out using a Nikon Eclipse Ti2 TIRF microscope with Hamamatsu Orca 4.0 sCMOS camera operated by Nikon Elements software. The TIRF angle was adjusted to pseudo-TIRF to create an evanescent field of approximately 300 nanometers, extending above the bottom of the pillars to ensure illumination of the membrane on the top of the pillars. Best illumination of the membrane and endocytic fluorophores was generated by aligning the illumination angle parallel to the direction of the nanoridges, as determined empirically.

Confocal microscopy for control siRNA and CLTC knockdown cells was done on a Zeiss LSM 900 with Airyscan 2.0 detection, in 4Y multiplex mode, operated by ZEN Blue software. Entire cell volumes were collected at Nyquist sampling frequency (140 nm per Z-section), and maximum intensity projections were created in ImageJ for analysis. Images were batch-processed by ZEN Blue software for Airyscan reconstructions using filter strength setting of 6.

Confocal microscopy for AP2 knockdown cells and for no-treatment transferrin uptake assays was done on a Nikon Eclipse Ti microscope with Yokogawa spinning disk and Andor EMCCD, operated by Nikon Elements software. Entire cell volumes were collected at Nyquist sampling frequency (200nm per Z-section), and maximum intensity projections were created in ImageJ for analysis.

### Computational Analysis

Puncta of endocytic proteins were detected using Trackmate 6.0.3 in Differences-of-Gaussians mode. Nanoridges were detected using custom Python scripts built on the Hough transform in Python package cv2. TIRF videos were analyzed using the MATLAB cmeAnalysis package. Standard settings for model-based detection, Gaussian fitting, radius of detection, and gap closing were selected. Further postprocessing was completed using custom Python scripts, available at github.com/drubinbarneslab. Valid endocytic sites were selected based on lifetime limits of 18-180 seconds, MSD measurement of 0.02 micron^2, and at least 4 consecutive seconds of dynamin2 detection, as have been previously reported for valid CCPs (Pascolutti et al., 2019).

Statistical analyses were conducted either with python packages NumPy, Pandas, SciPy, and Seaborns, or with Prism 9. Graphs were produced with Prism 9.

## Acknowledgments

We gratefully acknowledge Danielle Jorgens, Reena Zalpuri, and Guangwei Min of the UC Berkeley Electron Microscopy Lab for consultation and assistance in TEM and SEM imaging. Spinning-disk confocal imaging was conducted at the CRL Molecular Imaging Center, supported by the Gordon and Betty Moore Foundation. We also thank Stefano Cabrini, Scott Dhuey, Giuessepe Calafiore, and Stefano Dallorto of the Lawrence Berkeley Lab Molecular Foundry for assistance in silicon mold EBL/etching and Ormocomp substrate manufacturing, and Paul Lum of the UC Berkeley Biomoleuclar Nanotechnology Center for advice and assistance in Ormocomp lithography. Work at the Molecular Foundry was supported by the Office of Science, Office of Basic Energy Sciences, of the U.S. Department of Energy under Contract No. DE- AC02-05CH11231. We thank James Hurley and Leanna Owen of UC Berkeley and Ross Pedersen of the Carnegie Institute for comments on the manuscript. The authors declare no competing financial interests.

## Funding

National Institutes of Health grant R35GM118149 (DGD)

## Author contributions

Experiments were conceptualized by RCC and DGD, and carried out by RCC. Data analysis and visualization were done by RCC and CRS. RCC and DGD wrote the manuscript; all authors contributed to editing.

---

**Fig S1.**
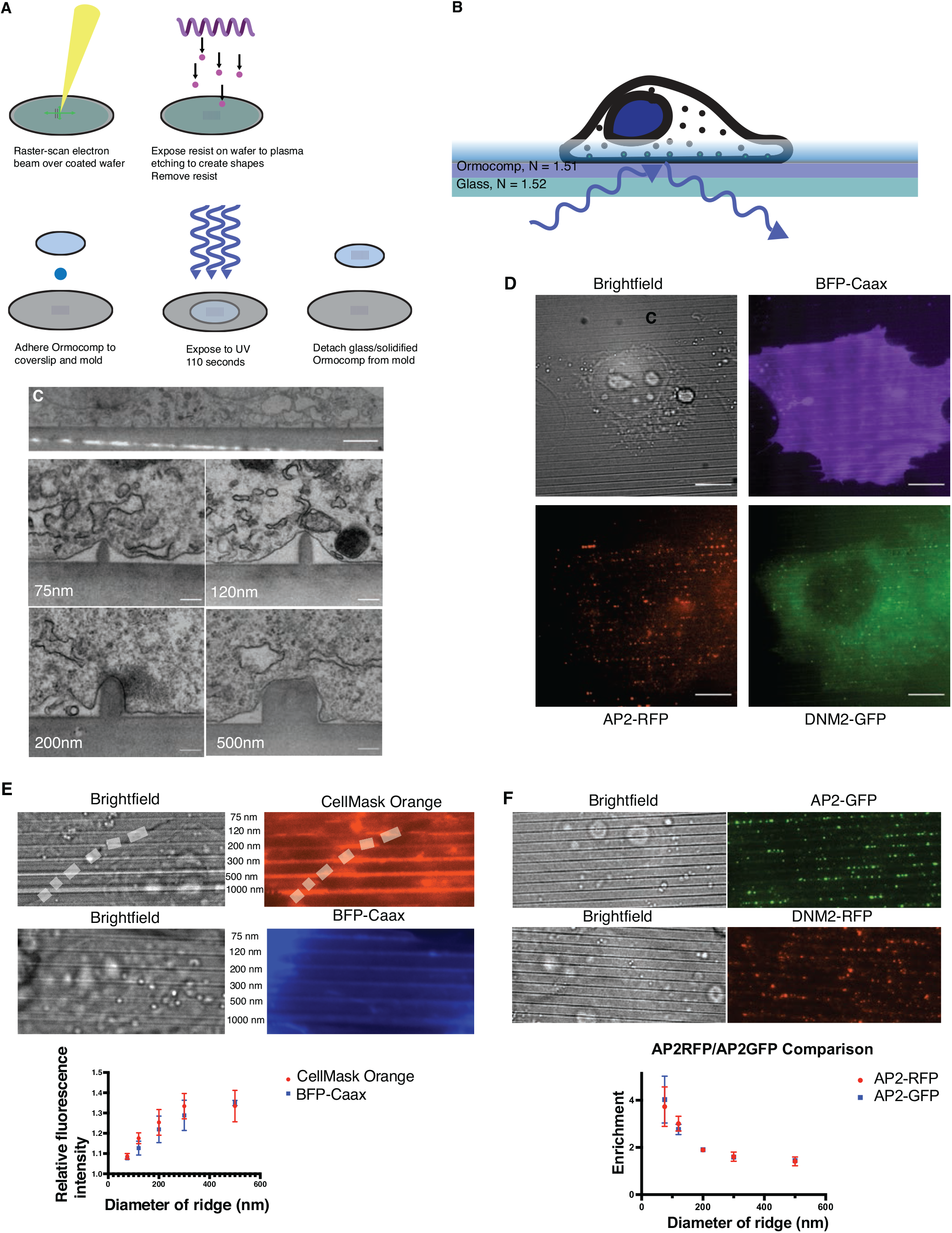
**(A)** Method of electron-beam lithography/reactive ion etching to produce the mold (top) and UV-nanoimprint lithography (bottom), to produce Ormocomp substrates on a microscope coverslip. **(B)** TIRF microscopy is possible on Ormocomp substrates because of the near-exact refractive index match of Ormocomp with glass. **(C)** Representative electron micrographs from Ormocomp nanoridges inducing semi-circular membrane shapes similar to the high-energy U-shaped intermediate of CME. Upper scale bar: 2 µm. Lower scale bar: 100 nm. **(D)** Brightfield image and 3-color TIRF microscopy of a cell on 120nm nanoridges with BFP-Caax marking the membrane, as well as AP2RFP and DNM2GFP preferentially localizing to the sites of curvature. Scale bar: 10 µm. **(E)** Cellmask Orange and BFP-Caax membrane staining showing an increase in membrane fluorescence on sites of high curvature, with quantification used in correcting for excess membrane area. Mean +/- SEM. N ≥ 3 cells per condition. White dashed line marks edge between two cells. **(F)** AP2GFP and DNM2RFP on 75nm nanoridges, showing that swapped fluorophore does not affect curvature enrichment of endocytic proteins. Mean +/- SD. N ≥ 3 cells per condition.

**Fig S2.**
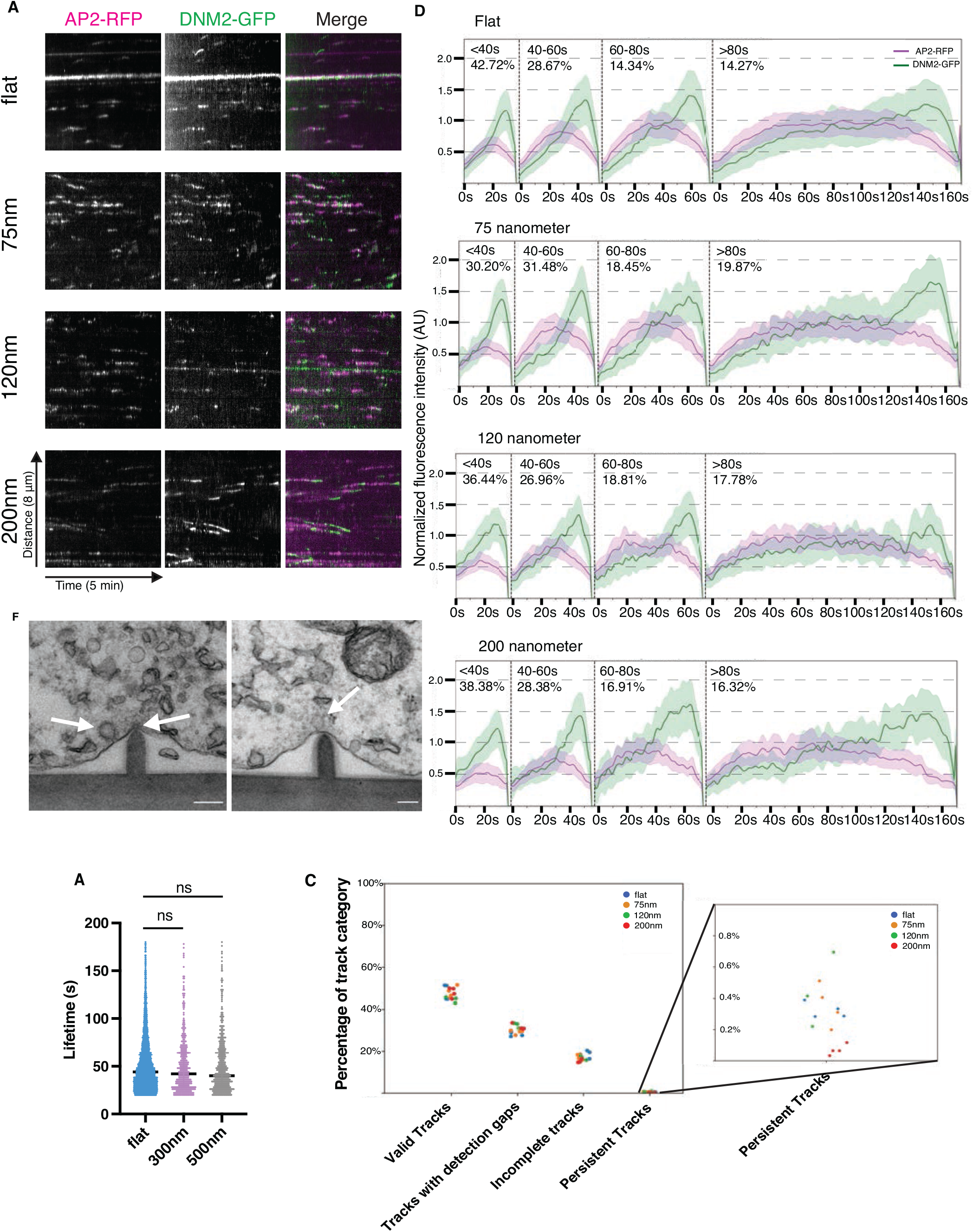
**(A)** Kymographs from two-color AP2RFP/DNM2GFP double-labeled cells from flat and curved substrates showing fluorescence intensity profiles indicative of productive endocytosis. **(B)** Fluorescence intensity profiles of endocytic sites, binned by lifetime, from flat and curved substrates showing profile of endocytic sites. N ≥ 1,000 tracks per condition. **(C)** TEM images of cells on 120nm nanoridges with white arrows indicating CCPs at early (on ridge) and late (left of ridge) stages. Scale bar: 200 nm. **(D)** Average lifetimes of endocytic sites from flat, 300nm, and 500nm substrates indicating no lifetime change on ridge sizes greater than 200 nm. P-value > 0.05, Student’s T test. Mean with interquartile range. N ≥ 300 tracks per condition. **(E)** Proportion of valid, gapped, incomplete, and persistent tracks on flat and curved substrates indicating no difference in proportion as a function of curvature. Each point is one cell, N ≥ 3 cells per condition.

**Fig S3.**
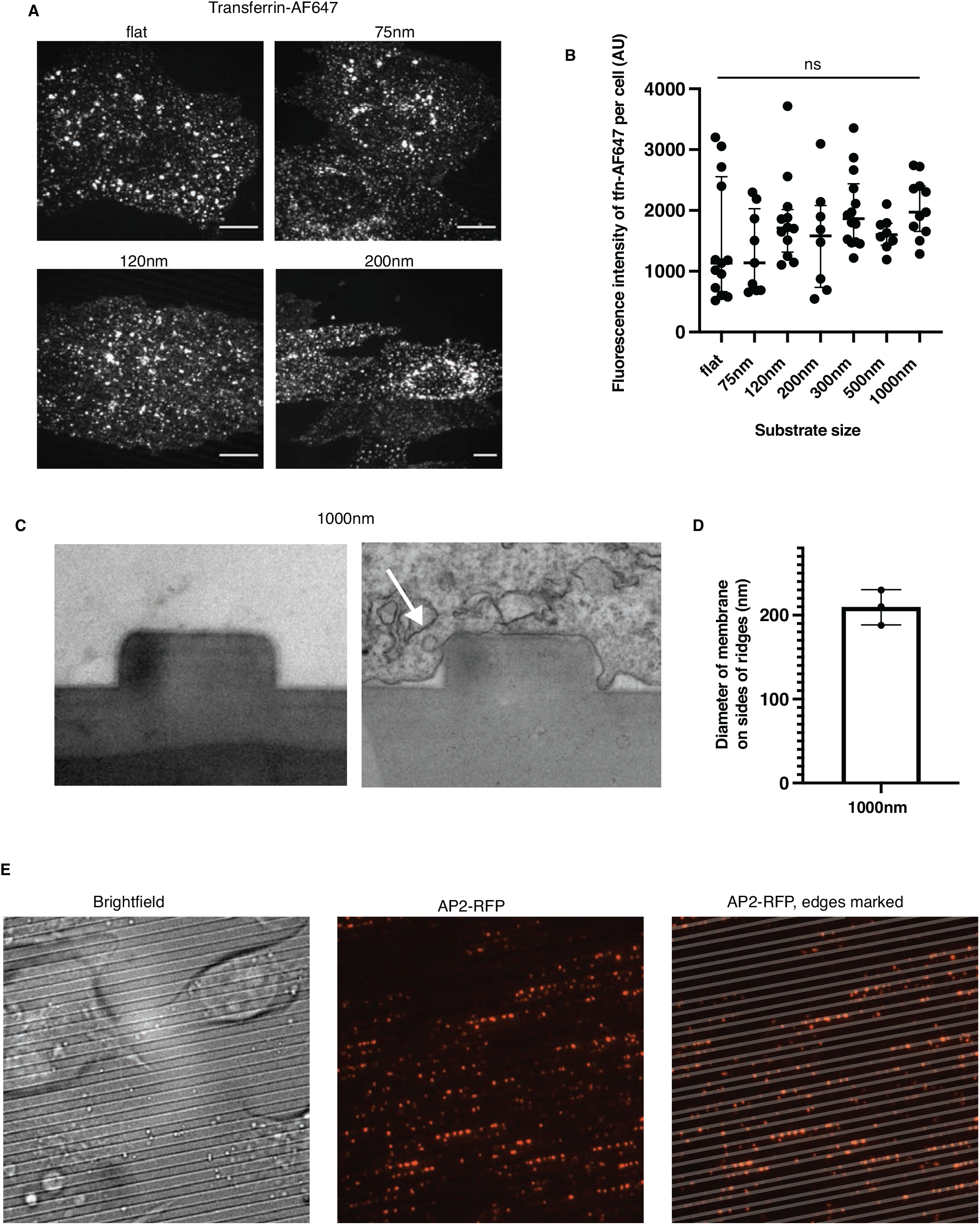
**(A)** Representative images of AlexaFluor647-tagged transferrin (tfn-AF647) from cells on flat and curved substrates. Scale bar: 10 µm. **(B)** Quantification of mean pixel intensity of tfn- AF647 from 10 cells plated on flat and curved substrates. Mean +/- SD. P-value > 0.05, Two-way ANOVA test. N = 10 cells per condition. **(C)** TEM images of bare 1000nm substrate and a cell grown on a 1000nm substrate showing induced curvature on the sides, but not the top, of the substrate. White arrow indicates a CCP growing from the high-curvature side. **(D)** Quantification of the edge membrane diameter from cells grown on 1000nm substrate. Mean +/- SD. N = 3 ridges from 2 cells. **(E)** Brightfield, AP2RFP, and merge images of cells on the 1000nm substrate demonstrating preferential AP2RFP localization on the edge of the ridge, rather than the top.

**Fig S4.**
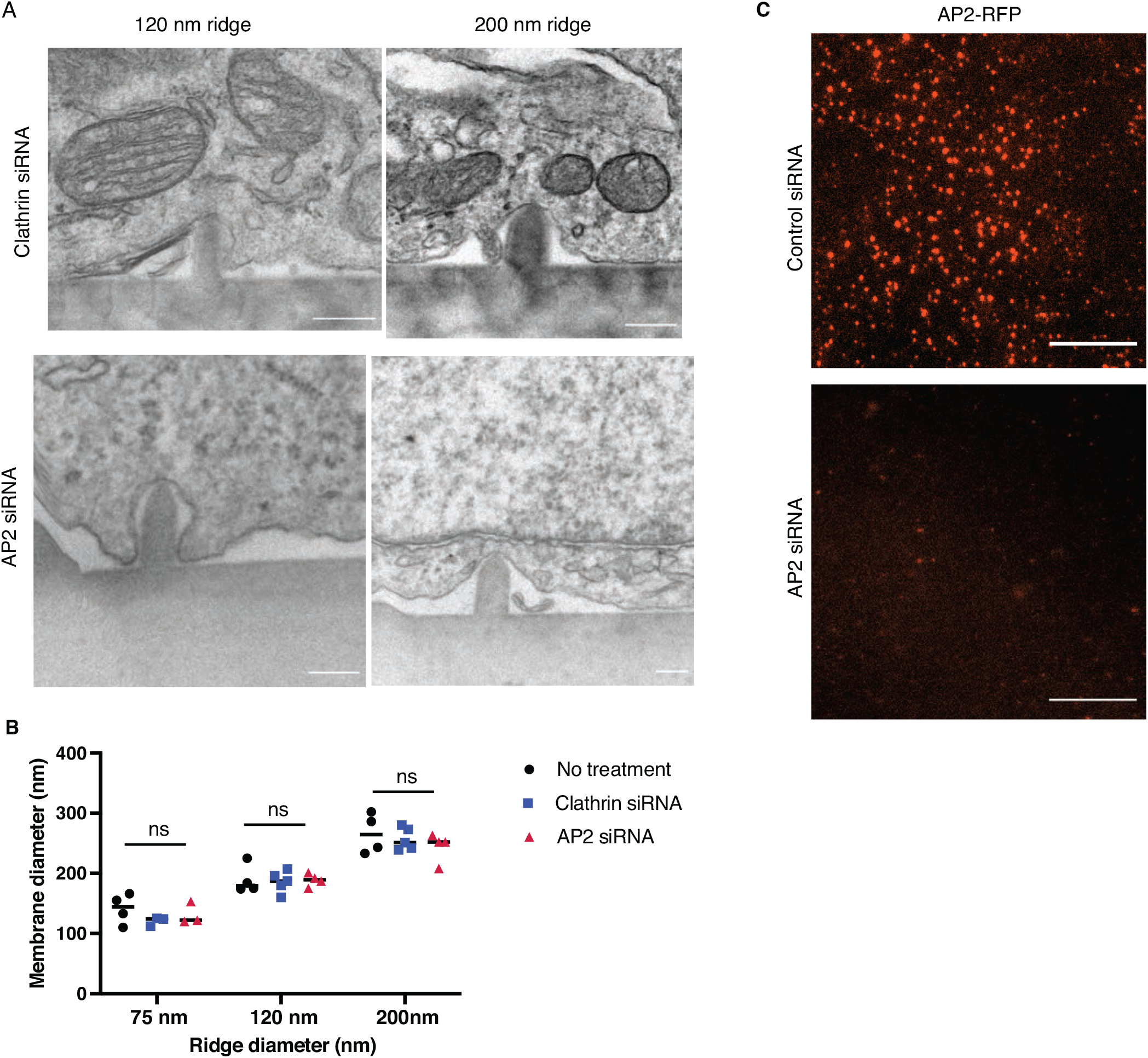
**(A)** Representative TEM images of clathrin knockdown or AP2 knockdown cells showing similar induced curvature to no-treatment cells. Scale bar: 200 nm. **(B)** Diameter induced by nanoridges measured from TEM images. Mean +/- SD. P-value > 0.05, two-way ANOVA test. N = 3 ridges from 3 cells per condition. **(C)** AP2RFP cells with control or AP2 siRNA showing the reduction in fluorescence intensity upon AP2 knockdown.

## Notes

### Competing Interest Statement

The authors have declared no competing interest.

### Summary of Updates

Additional references have been added; additional wording has been added for context of the findings. No findings have been changed.

